# Slipped strand mispairing in the gene encoding sialidase NanH3 in *Gardnerella* spp

**DOI:** 10.1101/2020.09.16.300947

**Authors:** Shakya P. Kurukulasuriya, Mo H. Patterson, Janet E. Hill

## Abstract

Cell wall proteins with sialidase activity are involved in carbohydrate assimilation, adhesion to mucosal surfaces, and biofilm formation. *Gardnerella* spp. inhabit the human vaginal microbiome and encode up to three sialidase enzymes, two of which are suspected to be cell wall associated. Here we demonstrate that the gene encoding extracellular sialidase NanH3 is found almost exclusively in *G. piotii* and closely related *Gardnerella* genome sp. 3, and its presence correlates with sialidase positive phenotype in a collection of 112 *Gardnerella* isolates. The *nanH3* gene sequence includes a homopolymeric repeat of cytosines that varies in length within cell populations, indicating that this gene is subject to slipped-strand mispairing, a mechanisms of phase variation in bacteria. Variation in the length of the homopolymer sequence results in encoding of either the full length sialidase protein or truncated peptides lacking the sialidase domain due to introduction of reading-frame shifts and premature stop codons. Phase variation in NanH3 may be involved in immune evasion or modulation of adhesion to host epithelial cells, and formation of biofilms characteristic of the vaginal dysbiosis known as bacterial vaginosis.

## Introduction

Bacterial vaginosis (BV) is a condition that is characterized by altered composition of the vaginal microbiota and occurs when the healthy microbiota is replaced by an overgrowth of mixed aerobic and anaerobic species, including *Gardnerella* spp. (1, 2). An abundance of *Gardnerella* spp. is often found in cases of symptomatic BV, although they are also found in healthy women with no clinical signs or symptoms of BV (3). *Gardnerella* spp. can be resolved into four subgroups based on cpn60 barcodes sequencing (4) or whole genome sequencing (5). Recently, the description of *Gardnerella vaginalis* was amended and three new species were defined within the genus *Gardnerella: G. leopoldii, G. swidsinskii*, and *G. piotii* (6). *G. piotii* and *G. vaginalis* correspond to cpn60 subgroup B and C, respectively. *G. leopoldii* and *G. swidsinskii* were previously grouped together as subgroup A based on cpn60 sequences of the isolates available at that time. Isolates belonging to cpn60 subgroup D corresponded to three distinct genome species: genome species 8, 9 and 10 (7).

In addition to the characteristic change in microbiota, and elevated pH, sialidase activity in vaginal fluid is a diagnostic marker of BV (8, 9). Sialidase enzymes cleave the glycosidic linkages of sialic acids from terminal glycans including vaginal mucins, immunoglobulin A molecules, and epithelial cell surface glycoproteins (10, 11). Activity on the latter may be related to the adhesion of bacteria to epithelial cells; the initial step in biofilm formation (12). Previously, we have demonstrated that sialidase activity is almost exclusively confined to *G. piotii* and *Gardnerella* genome species 3 (cpn60 subgroup B) (13).

A putative sialidase gene, *nanH1* (sialidase A) was identified in *Gardnerella* spp. and was initially thought to be the gene responsible for sialidase activity (14). Although *nanH1* appears to be present in all sialidase positive strains, it is also found in sialidase activity negative strains (13). This observation combined with the lack of signal peptide on the NanH1 proteins suggests that this protein is an intracellular enzyme, likely involved in manipulation of nutritional substrates as is the case in many bacteria (15). The discrepancy between *nanH1* presence and enzyme activity led to the recognition of two additional sialidase genes (*nanH2* and *nanH3*) in some *Gardnerella* isolates (16), which are unevenly distributed among *Gardnerella* spp. (17). Based on biochemical studies, Robinson et al. (16) concluded that NanH2 and NanH3 are the proteins responsible for sialidase activity detected in vaginal fluid. The presence of signal peptides and/or C-terminal transmembrane alpha-helices in some of the identified sequences further suggested that they are secreted and tethered to the cell, although some proportion may subsequently be released into the extracellular environment due to proteolytic activity (16). Cell wall anchored proteins with sialidase activity in other species have been found to be involved in adhesion to mucosal surfaces and biofilm initiation as well as carbohydrate assimilation (18–21).

Interestingly, in our examination of genome sequences of our collection of *Gardnerella* isolates, we noted that *nanH3* contains a homopolymeric tract of cytosine residues. Genomic regions that contain short, homogenous or heterogenous repeats are susceptible to slipped-strand mispairing in which the length of the repeat region can change with each replication (22). The result of this modulation is phase variation: a reversible process in which the expression of the encoded protein can be rapidly switched on and off (23). Phase variation in cell surface proteins can result in immune evasion and alteration of biofilm phenotypes (24, 25).

Here we determined the distribution of genes encoding sialidases NanH2 and NanH3 in the context of newly reclassified *Gardnerella* spp. and demonstrated that *nanH3* is subject to slipped-strand mispairing.

## Methods

### Protein domain identification and sequence alignment

DNA and protein sequence alignments were performed with Clustal Omega (EMBL_EBI) and NCBI BLAST (basic local alignment search tool). InterproScan (https://www.ebi.ac.uk/interpro/) and SignalP were used to predict the location of functional domains and signal peptides (26).

### Bacterial strains and culture conditions

*Gardnerella* strains (n = 112) from a previously described culture collection were used in the study (13). Sialidase activity for all isolates had been determined previously (13) using a quantitative assay and fluorogenic substrate 2’-(4-methylumbelliferyl)-α-D-N-acetylneuraminic acid sodium salt hydrate (11), and whole genome sequences for 36 of these isolates were previously determined (27). Complete strain information and sequence accessions are provided in Table S1.

*Gardnerella* isolates were grown on Columbia Sheep Blood Agar plates (BBL, Becton, Dickinson and Company, Sparks, MD, USA) at 37°C for 48 hours with anaerobic BD GasPak EZ (Becton, Dickinson and Company, Sparks, MD, USA). For broth cultures, a few colonies from the plate were collected with a 10 μl inoculation loop and used to inoculate NYC III (ATCC 1685 medium; per litre: 2.4g HEPES, 15 g Proteose peptone, 3.8 g Yeast extract, 5 g NaCl, 5 g Glucose). Broth cultures were incubated at 37°C for 48 hours in anaerobic conditions.

### PCR screen for *nanH3*

Genomic DNA was purified from broth cultures using a modified salting-out procedure (28). All DNA extracts were initially tested by PCR for the universal cpn60 barcode to confirm the quality of the DNA.

To screen isolates for the presence of *nanH3*, PCR primers were designed based on multiple sequence alignments of 15 *nanH3* sequences obtained from the Integrated Microbial Genomes database (https://img.jgi.doe.gov/). Degenerate primers were designed to account for sequence variability within the gene sequence and to amplify a product of 375 bp in length (JH0684: 5’-GTT GTA GAR CTT TCT GAT GG-3’, JH0685: 5’-YRY TAT TAT CGC CCT CAT ATA-3’). PCR reactions contained 1 × PCR Buffer (0.2 M Tris-HCl at pH 8.4, 0.5 M KCl), 2.5 μM MgCl_2_, 0.40 μM dNTP, 0.20 μM forward primer, 0.20 μM reverse primer, 2 U Taq DNA Polymerase, ultrapure water and 2 μl of template DNA in a final volume of 50 μl. PCR reactions were conducted using the following thermocycling parameters in a Mastercycler Pro 6321 (Eppendorf AG, Hamburg, Germany): 94 °C for 3 minutes, 40 cycles of (94 °C for 30 seconds, 55 °C for 30 seconds, 72 °C for 30 seconds), 72 °C for 1 minute, hold at 20 °C. PCR products were visualized under UV light on a 1.0 % (w/v) agarose gel containing ethidium bromide.

### Homopolymer PCR, cloning and sequencing

To determine the length of the homopolymer region of *nanH3*, four strains (W11, VN014, VN015, NR032) were cultured on Columbia Agar plates with 5% (v/v) sheep blood. Primers JH0780 (5’-ATG ATT GGA ACA GCG CAT AAA G-3’) and JH0781 (5’-GAT TTC TCC ACC TAC AGT TAC C-3’) were designed to PCR amplify a region including basepairs 2-310 of the open reading frame of *nanH3*.

The DNA sequence (308 bp) was amplified by PCR in a Mastercycler Pro 6321 (Eppendorf AG, Hamburg, Germany). The components of the PCR reaction mix (50 μl per reaction) were added to achieve final concentrations of 1×High Fidelity PCR buffer (60 mM Tris-SO_4_ (pH 8.9), 18 mM (NH_4_)_2_SO_4_), 0.2 mM dNTP mix, 2 mM MgSO_4_, 1U Platinum High Fidelity (Hi-Fi) proof reading Taq polymerase (Invitrogen, Carlsbad, CA, USA). Two colonies of each *G. piotii* or genome sp. 3 strain were randomly picked and added to separate PCR reactions using sterile toothpicks. Thermocycling conditions included 35 cycles of denaturation at 94°C for 15 seconds, annealing at 55°C for 30 seconds, extension at 68°C for 1 minute, and final extension at 68°C for 5 minutes. PCR products were visualized on a 1% (w/v) agarose gel. PCR products were purified using QiaQuick PCR purification kit (Qiagen, Hilden, Germany) and purified PCR products were sequenced using the amplification primers (JH0780, JH0781).

To clarify the exact length of the homopolymer region, amplicons generated from the two colonies of each of the four strains were A-tailed in 10 μl reactions containing 1× Platinum PCR buffer (20 mM Tris HCl (pH 8.4), 50 mM KCl), 2 mM MgCl_2_, 0.5 mM dATP, 5 U Platinum Taq polymerase (Invitrogen, Carlsbad, CA, USA) and less than 500 ng of the purified PCR product. The reaction mixture was incubated at 72 °C for 20 minutes in Mastercycler Pro 6321 (Eppendorf AG, Hamburg, Germany). End-modified PCR products were ligated into pGEM-T Easy vector (Promega, Madison, WI, USA). The vector-insert construct was used to transform chemically competent DH5α cells or OneShot Top 10 *E. coli* (Invitrogen, Carlsbad, CA, USA) and plated on LB/AMP/X-gal agar media. Ten white colonies were randomly selected from the transformants of each *Gardnerella* strain and transferred into LB + ampicillin broth. Cultures were grown overnight at 37 °C. Plasmid DNA was isolated using QiaPrep Spin Miniprep kit (Qiagen, Hilden, Germany). Plasmids were sequenced using vector primers T7 and SP6.

## Results

### Distribution of extracellular sialidases in *Gardnerella* spp. isolates

Robinson et al. (16) had previously reported that sialidase activity was associated with the presence of either *nanH2* or *nanH3* in a collection of 34 *Gardnerella* isolates but did not report the species or subgroup affiliation. In order to confirm this observation and to reconcile the distribution of genes and activities with the new *Gardnerella* taxonomic framework (6), we queried an unrelated set of 36 isolates in our culture collection for which whole genome sequence data and sialidase activity data were available (Table S1). The presence of the genes was assessed by aligning *nanH2* of *G. piotii* JCP8151B (ATJH01000056) and *nanH3* of *Gardnerella* genome sp. 3 strain W11 to the genome sequences using BLASTn. BLASTn results were dichotomous, with either one clear match per genome (E = 0.0, bitscore 1275-5049, percent identity 95-99%) or no significant hits reported. Sialidase activity was reported in 13/36 isolates (Table 1) including 12 *G. piotii* or genome sp. 3 (cpn60 subgroup B) isolates and one *G. vaginalis* isolate. The *nanH3* gene was found in 12/13 sialidase activity positive isolates while *nanH2* was present in only 7/13 isolates. Six isolates possessed both *nanH2* and *nanH3* genes in their genomes. VN002 was sialidase activity positive but only *nanH2* was identified in the genome sequence. This isolate subsequently screen PCR positive for *nanH3* (see next section), suggesting that the gene was not included in the shotgun assembly of the genome, which consisted of multiple gap-containing scaffolds. None of the *G. swidsinskii, G. leopoldii* or subgroup D isolates were sialidase positive and none contained *nanH2* or *nanH3*.

**Table 1.** Distribution of *nanH2* and *nanH3* genes in *Gardnerella* spp. whole genome sequences

| Isolate | cpn60 Subgroup | Species | Sialidase activity<sup>1</sup> | *nanH2* | *nanH3* |
| --- | --- | --- | --- | --- | --- |
| GH007 | B | *G. piovii* | + | + | + |
| GH019 |  | *G. piovii* | + | + | + |
| GH020 |  | *G. piovii* | + | + | + |
| GH022 |  | *G. piovii* | + | + | + |
| N170 |  | Genome species 3 | + | - | + |
| NR026 |  | Genome species 3 | + | - | + |
| VN002 |  | *G. piovii* | + | + | -<sup>2</sup> |
| N144 |  | Genome species 3 | + | - | + |
| W11 |  | Genome species 3 | + | - | + |
| N101 |  | Genome species 3 | + | + | + |
| N153 |  | Genome species 3 | + | + | + |
| N95 |  | Genome species 3 | + | - | + |
| GH005 | A | *G. leopoldii* | - | - | - |
| NR015 |  | Unknown | - | - | - |
| NR016 |  | *G. swidsinskii* | - | - | - |
| NR017 |  | *G. leopoldii* | - | - | - |
| NR019 |  | *G. leopoldii* | - | - | - |
| NR020 |  | *G. swidsinskii* | - | - | - |
| NR021 |  | *G. swidsinskii* | - | - | - |
| VN003 |  | *G. leopoldii* | - | - | - |
| WP021 |  | *G. swidsinskii* | - | - | - |
| WP022 |  | *G. swidsinskii* | - | - | - |
| NR010 |  | *G. leopoldii* | - | - | - |
| N072 |  | *G. swidsinskii* | - | - | - |
| GH021 | C | *G. vaginalis* | - | - | - |
| NR001 |  | *G. vaginalis* | - | - | - |
| NR037 |  | *G. vaginalis* | - | - | - |
| NR038 |  | *G. vaginalis* | - | - | - |
| NR039 |  | *G. vaginalis* | - | - | - |
| WP023 |  | *G. vaginalis* | - | - | - |
| N165 |  | *G. vaginalis* | - | - | - |
| GH015 |  | *G. vaginalis* | + | - | + |
| NR003 | D | Genome sp. 8 | - | - | - |
| NR047 |  | Unknown | - | - | - |
| WP012 |  | Genome sp. 9 | - | - | - |
| N160 |  | Genome sp. 10 | - | - | - |

<sup>1</sup>Previously determined (13)447 <sup>2</sup>PCR positive for *nanH3*

Interestingly, the NanH3 of *G. piotii* strain JCP8151B investigated by Robinson et al. was reported to lack a signal peptide (16). When we examined the sequence upstream of *nanH3* in the JCP8151B sequence (Genbank accession ATJH01000033), we found that there is an alternative start codon, and a cytosine homopolymer. The frame-shift that resulted in the annotation of the gene without the signal peptide-encoding N-terminus is caused by the homopolymer. This was also the case for isolates GH020, GH022, N170, and NR026 where the homopolymer created a frame-shift that resulted in a protein annotation lacking the signal peptide encoded by sequence immediately upstream of the homopolymer. In N144, W11, N101, N153 and N95, the homopolymer length did not create a frame-shift and the annotated NanH3 protein sequence includes a signal peptide. In GH007, GH019, and GH015 we were unable to determine if a signal-peptide encoding sequence was present due to assembly gaps immediately upstream of the homopolymer in these genomes (Table S1).

Signal peptides were detected in 4/7 predicted NanH2 sequences (GH019, GH020, N101 and N153). In the other three cases where a *nanH2* sequence was identified, we could not conclude if a signal peptide was included in the protein sequence provided in the Genbank annotation since assembly gaps in those regions of the genomes affected the annotation of these proteins.

### Correlation of *nanH3* with sialidase activity

In order to examine further the relationship of *nanH3* to sialidase activity, 112 *Gardnerella* spp. isolates for which sialidase activity data was available (13) were screened for the presence of *nanH3* using PCR primers JH0684/JH0685 (Figure 1). Sialidase activity was detected in 32/33 *G. piotii/genome* sp. 3 isolates and 3/35 *G. vaginalis* isolates. All sialidase activity positive isolates were positive for *nanH3* by PCR with the exception of one *G. piotii* isolate (WP027) that was PCR negative. Unfortunately, genome sequence data was not available for WP027.

**Figure 1.**
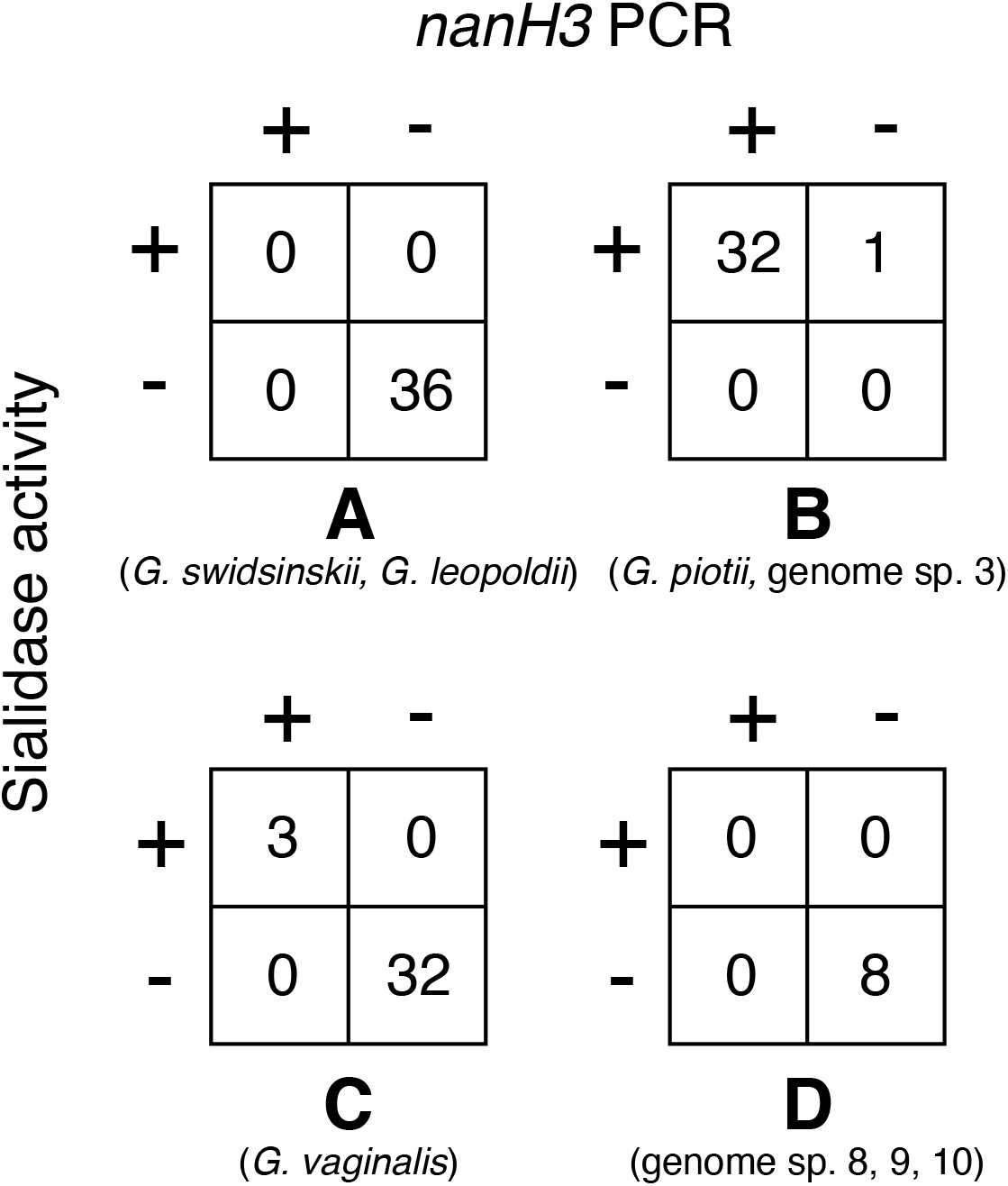
Correlation of *nanH3* and sialidase activity in 112 *Gardnerella* isolates. Numbers of isolates positive or negative for sialidases activity and *nanH3* PCR. cpn60 subgroup affiliation and species are indicated below each crosstab. Sialidase results were determined by Schellenberg *et al*. (13).

### Characterization of a poly-C homopolymer in *nanH3*

The protein sequences encoded by all nine *nanH3* sequences without assembly gaps were predicted to encode a signal peptide (amino acids 1-32), a sialidase domain (amino acids 183-547 in W11) and a C-terminal transmembrane domain (amino acids 789-807 in W11) (Figure 2A). We observed a poly-cytosine homopolymer (8-14 cytosines) in all *nanH3* genes identified in the whole genome sequences, approximately 100 bases from the start of the open reading frame. The homopolymer occurred immediately following the region of the sequence predicted to encode the signal peptide (amino acids 1-32) (Figure 2B). When Sanger sequencing was performed on PCR products corresponding to nucleotides 2-310 of *nanH3* amplified from genomic DNA extracted from broth cultures of strain W11, the results suggested variable lengths of the homopolymer. Specifically, clean data was obtained upstream of the poly-C tract but sequence 3’ to that region was indicative of a mixed template (Figure S1). Since it has been demonstrated that simple repeats such as homopolymers can be subject to slipped strand mispairing, we set out to determine if the homopolymer region of *nanH3* varied in length within and between strains of *G. piotii*/genome species 3.

**Figure 2.**
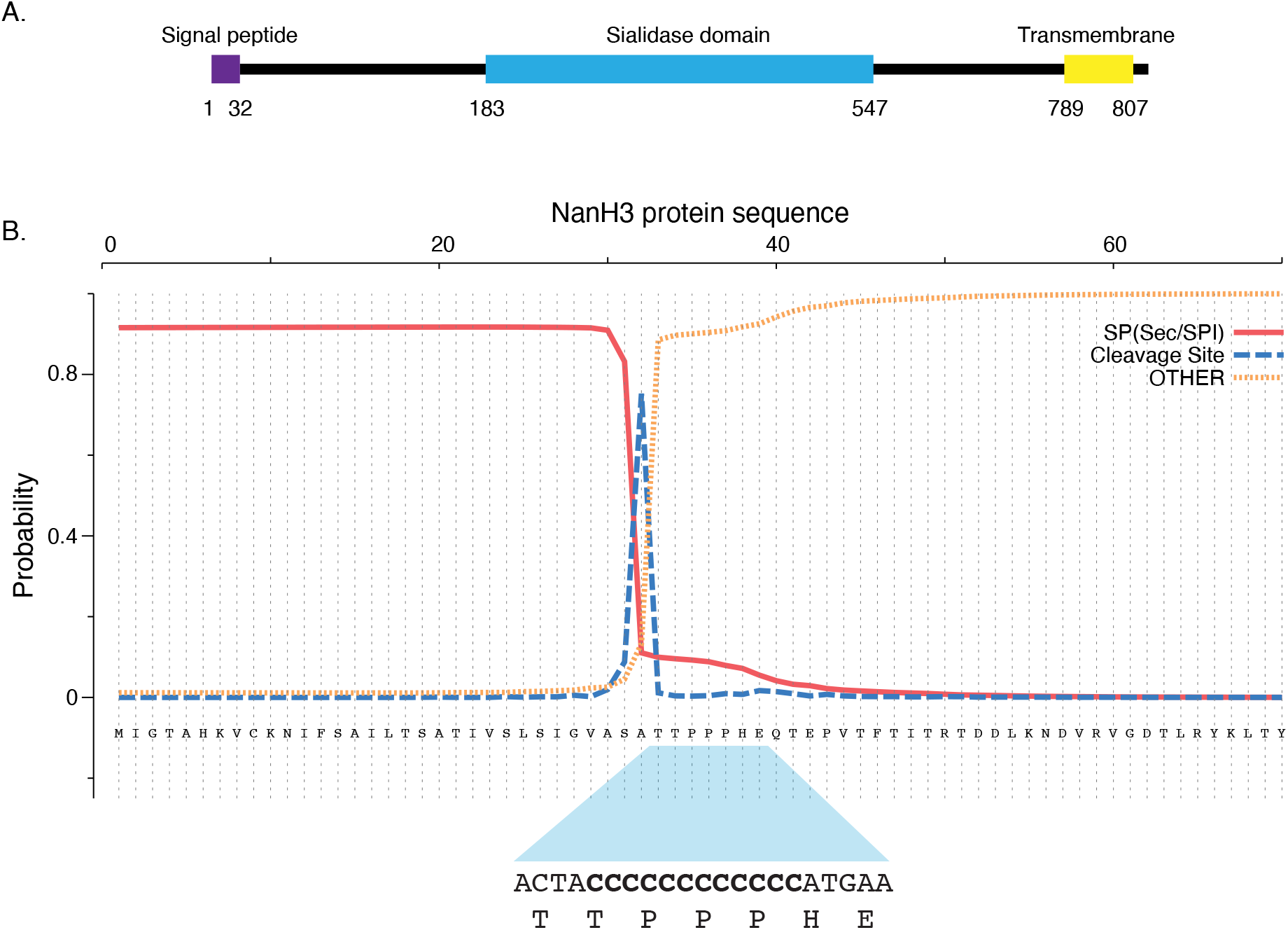
A. Predicted locations of signal peptide, sialidase and transmembrane domains of NanH3 from strain W11 based on InterProScan analysis. B. Location of predicted signal peptide, cleavage site and adjacent homopolymer in the *nanH3* gene of strain W11. Analysis was conducted with SignalP-5.0.

PCR product libraries were made from two colonies each of isolates W11, VN014, VN015 and NR032 (8 clone libraries total). Plasmids were purified from 10 colonies from each of the 8 clone libraries and sequenced. High quality sequence data was obtained from 15, 14, 18 and 16 clones from the W11, VN014, VN015 and NR032 libraries, respectively. Example results are shown in Figure S2. The length of the homopolymer varied from 8 to 14 among all strains (Table 2), and within each strain the sequence flanking the homopolymer was identical. *In silico* translation of the encoded polypeptides showed that most homopolymer lengths resulted in a truncated peptide (37-44 amino acids), while full-length protein (812 aa) would result when there were nine or twelve cytosines in the homopolymer region. Truncated peptides would include the signal peptide (amino acids 1-32) but no part of the predicted sialidase domain (amino acids 183-547) (Figure 2). Interestingly, the most common length of the poly-C region varied among strains. C_10_ was most common in VN014 and VN015, while C_9_ and C_12_ were most frequently observed in NR032 and W11, respectively. VN015 had the highest number of homopolymer variants (Figure 3). A second open reading frame within the *nanH3* open reading frame was also identified, corresponding to amino acids Met_397_-Tyr_812_ of the NanH3 protein. These 415 aa polypeptides encompass part of the predicted sialidase domain, and the C-terminal transmembrane domain but lacks a signal peptide.

**Table 2.** Homopolymer length variants in *Gardnerella piotii* and genome sp. 3 isolates.

| Isolate | Homopolymer sequence | Predicted peptide length (amino acids)<sup>1</sup> |
| --- | --- | --- |
| W11 | CAACTA- C<sub>11</sub>-ATGAACAAA | 43 |
|  | CAACTA- C<sub>12</sub>-ATGAACAAA | 812 |
|  | CAACTA- C<sub>13</sub>-ATGAACAAA | 38 |
|  | CAACTA- C<sub>14</sub>-ATGAACAAA | 44 |
| VN014 | CAACTA- C<sub>9</sub>-TCGAACAAA | 812 |
|  | CAACTA- C<sub>10</sub>-TCGAACAAA | 37 |
|  | CAACTA- C<sub>11</sub>-TCGAACAAA | 43 |
| VN015 | CAACTA- C<sub>9</sub>-TCGAACAAA | 812 |
|  | CAACTA- C<sub>10</sub>-TCGAACAAA | 37 |
|  | CAACTA- C<sub>11</sub>-TCGAACAAA | 43 |
|  | CAACTA- C<sub>12</sub>-TCGAACAAA | 812 |
|  | CAACTA- C<sub>13</sub>- -GAACAAA | 43 |
| NR032 | CAACTA- C<sub>8</sub>-ATGAACAAA | 42 |
|  | CAACTA- C<sub>9</sub>-ATGAACAAA | 812 |

<sup>1</sup>Full-length NanH3 is 812 amino acids

**Figure 3.**
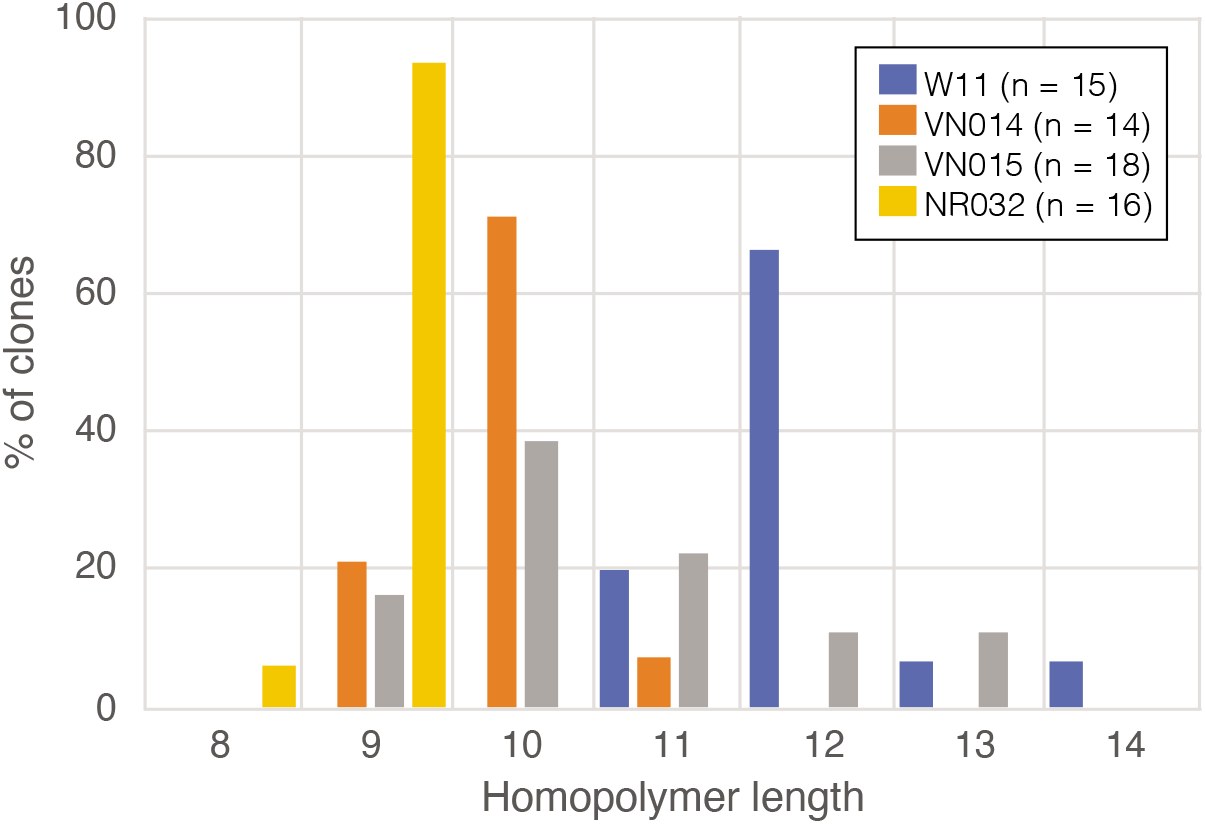
Frequencies of the homopolymer length variants in isolates W11, VN014, VN015 and NR032.

## Discussion

The genus *Gardnerella* is known as a hallmark of BV and their abundance is used as a criterion for the laboratory diagnosis of the condition (29). In addition to participating in biofilms that coat epithelial cells in BV, *Gardnerella* spp. produce enzymes and a cholesterol dependent cytolysin (vaginolysin) that contribute to degrading the protective barriers of the vaginal mucosa (30). Sialidase activity can provide nutrients by releasing sialic acid moieties from vaginal sialylated mucins (11), altering the physical properties of vaginal mucus. Removal of sialic acid residues from epithelial cell surface glycans can facilitate bacterial adhesion and initiation of biofilm formation (12, 18, 20, 21).

The two putative sialidase genes (*nanH2* and *nanH3*) in *Gardnerella* spp. have been demonstrated to encode proteins with high enzymatic potency (16). Taken together, the result of this study and our current study show that *nanH3* is more common and that *nanH2* is virtually never found without *nanH3.* It is now clear that extracellular sialidase activity is a property of *G. piotii* and the closely related *Gardnerella* genome sp. 3. We found three occurrences of sialidase positive, *nanH3* positive *G. vaginalis* isolates (3/35 isolates screened), and Vaneechoutte et al. (6) reported sialidase activity in 1/4 *G. vaginalis* isolates used in the amendment of the genus. This infrequent prevalence of *nanH3* in *G. vaginalis* could be the result of lateral gene transfer from *G. piotii* since these species co-exist in the same microbiome and women are usually colonized by more than one species (7, 31, 32).

The *nanH3* sequences we examined were predicted to encode a signal peptide, a sialidase domain and a C-terminal membrane domain, suggesting a cell-wall tethered protein, similar to SiaBb2 of *Bifidobacterium bifidum*, which enhances adhesion to intestinal mucosal surfaces and contributes to carbohydrate assimilation (21). The presence of the homopolymer immediately downstream of the signal peptide encoding region presents a challenge to automated annotation of this sequence. Sequencing and assembly methods commonly used in bacterial whole genome sequencing struggle with homopolymeric sequences, which can result in frame-shifts such as we observed in the *nanH3* sequence of JCP8151B.

Genomic regions that contain homogenous or heterogenous repeats are prone to changes in length of the repeat at each replication due to slipped strand mispairing (22). This can lead to consequent changes in the transcription or translation product of a gene depending on whether the slipped strand mispairing occurs within the open reading frame or in extragenic regions such as promoter sequences. Phase variation results when slipped strand mispairing creates “on” and “off” states of expression of a protein. Phase variation of cell surface proteins has been documented in many bacterial species including *Neisseria* spp. (33–35), *Salmonella* spp. (36–38), *Treponema pallidum* (39, 40) and *Helicobacter pylori* (25). It was first described in the *opa* genes that encode opacity surface proteins of *Neisseria* spp. (41). The *opa* genes contain CTCTT pentamer repeats within the signal peptide encoding region and the expression of the Opa protein is regulated by slipped-strand mispairing. With six, nine or twelve CTCTT repeats the initiation codon is in frame with the remaining *opa* gene translating the Opa protein. Four or eight coding repeats makes the initiation codon out of frame with rest of the *opa* codons. The phase “on” state allows them to adhere to specific surface receptors on host cells (42).

Our results clearly show variation of the homopolymer length among *G. piotii* and *Gardnerella* genome sp. 3 within colonies growing on agar, with anywhere from 8 to 14 C’s observed. The effect of this change would be a mixture of cells with translation of NanH3 on or off. Why would *G. piotii* have a sialidase subject to phase variation? Two possible reasons are immune evasion and cell adhesion (biofilm initiation and dispersal).

Although BV is often referred to as a non-inflammatory condition because of the lack of typical clinical signs of inflammation (leukocyte infiltration, pain, redness, swelling), there is evidence of a host immune response to bacteria associated with BV, including *Gardnerella* spp.. IgA specific for *Gardnerella* produced vaginolysin has been detected in women with clinical signs of BV (43), and IgA levels correlate with IL-8 expression in vaginal secretions (44). *Gardnerella* has also been shown to stimulate pro-inflammatory cytokines *in vitro* (45). Balancing this inflammatory response are the actions of bacterial enzymes like prolidases and sialidases that can degrade the effectors of inflammation. Whether there is a specific host response to NanH3 remains to be determined.

The initial step in biofilm formation is adhesion, which in the case of BV associated biofilms is to epithelial cells (2). *Streptococcus pneumoniae* surface proteins with sialidase activity have been shown to reveal carbohydrate ligands for bacterial adhesion to host cells (18). Similarly, Soong et al. (19) showed that a cell wall associated sialidase of *Pseudomonas aeruginosa* plays a critical role in facilitating respiratory mucosa infection through biofilm formation. Specific antibodies against *B. bifidum* SiaBb2, a cell wall tethered sialidase, inhibit adhesion to cells (21). The involvement of *Gardnerella* sialidases in adhesion has not been demonstrated directly, although it has been reported that inhibition of sialidase activity in *Gardnerella* strain JCP8066 by an antiviral drug (Zanamavir) resulted in reduction of adherence to vaginal epithelial cells *in vitro* (12). *Gardnerella* spp. can form multi-species biofilms *in vitro* (46) and although “*Gardnerella vaginalis*” has been shown to participate in multispecies biofilms with other BV associated bacteria *in vivo* (47, 48) and *in vitro* (49), it is not yet known how each of the *Gardnerella* spp. participate in this process. Interestingly, *G. piotii* and *Gardnerella* genome sp. 3 (cpn60 subgroup B) have been previously associated with “intermediate” grades of vaginal dysbiosis (7, 31, 50, 51) and may contribute to transition from eubyosis to dysbiosis and enhance colonization by other BV associated anaerobes. Phase variation in a cell surface associated sialidase enzyme might be critical for turning on or off the initial adhesion of *Gardnerella* to epithelial cells and thus the cascade of events following that result in the biofilm covered “clue cells” typical of BV (52).

It seems likely that some of the NanH3 protein produced by *G. piotii* does not remain tethered to the cell surface, as is the case with *S. pneumoniae* (53), since the sequence upstream of the C-terminal transmembrane domain may be susceptible to proteolytic cleavage, and heterologously expressed NanH3 lacking the transmembrane domain is active (16). Taken together, the activity of NanH3 while anchored to the cell surface or released into the extracellular environment, and the likelihood that it is subject to phase variation through slipped-strand mispairing, make this protein an intriguing puzzle for future study. Development of genetic tools to create mutants where NanH3 production is locked “on” or “off” would be a significant step toward understanding the role of this protein in the initiation and maintenance of vaginal dysbiosis.

## Acknowledgements

This research was funded by a Natural Sciences and Engineering Council of Canada Discovery Grant to JEH. The authors are grateful to Champika Fernando for excellent technical assistance, and to all our colleagues in the Hill Lab for helpful discussions and feedback.

## Supplemental Materials

**Table S1.** *Gardnerella* strains used in the study, cpn60 subgroup affiliations, whole genome sequence accession numbers, sialidase activity phenotypes, presence/absence of *nanH2, nanH3*, signal peptide encoding sequences and homopolymers.

**Figure S1.** Sequencing of PCR products amplified directly from colonies of isolate W11.

PCR was performed with primers flanking the homopolymer region as described in the methods. In each electropherogram, after eleven peaks of cytosine, the sequencing signal shows multiple overlapping peaks suggesting the presence of multiple template sequences with different homopolymer lengths.

**Figure S2.** Electropherograms of the homopolymer regions in two W11 colonies. Colony 6 had individuals with 11, 12 and 13 C’s in the homopolymeric region. Five out of eight colonies had 12C’s in the homopolymer region. Colony 8 had individuals with 11, 12 and 14 C’s in the homopolymeric region. Five out of eight colonies had 12 C’s in the homopolymer region. Having 12 C’s makes the coding region in-frame. When there is 11,13 or 14 C’s, a premature STOP codon is generated possibly making the protein truncated.

